# Conformational fluctuations in molten globule state of *α*-lactalbumin

**DOI:** 10.1101/2022.05.13.491909

**Authors:** Abhik Ghosh Moulick, J. Chakrabarti

## Abstract

Molten globule (MG) state is an intermediate state of protein observed during folding into native structure. MG state of protein is induced by various denaturing agent (like Urea), extreme pH, pressure and heat. Experiments suggest that MG state of some protein is functionally relevant even if there is no well-defined tertiary structure. Earlier experimental and theoretical studies suggest that MG state of the protein is dynamic in nature, where conformational states are interconverted in nanosecond time scales. These observations lead us to study and compare conformational fluctuations of MG state to those of intrinsic disordered protein (IDP). We consider *α*-Lactalbumin(aLA) protein, which shows MG state at low pH upon removal of calcium (Ca^2+^) ion. We use constant pH molecular dynamics simulation (CpHMD) to maintain low pH during simulation. We use the dihedral principal component analysis, the density based clustering method and the machine learning technique to identify the conformational fluctuations. We observe metastable states in the MG state. The residues containing the essential coordinates responsible for metastability belong to stable helix in crystal structure, but most of them prefer unstructured or bend conformation in MG state. These residues control the exposure of the putative binding residues for fatty acids. Thus, the MG state of protein behaves as intrinsic disorder protein, although the disorder here is induced by external conditions.

## 1 Introduction

Some proteins show structural fluctuations in some parts in a near denaturing condition, while retaining their overall tertiary structures. Such states are called Molten Globule (MG) state of the protein.^1^ The MG state is induced by various denaturing conditions like high temperature, pH, high pressure and due to presence of various denaturing chemicals like urea.^2–4^ MG states, despite having structural fluctuations, show binding with ligands. Many of the complexes have functional relevance. For instance, the complexes of fatty acids and MG of *α*-lactalbumin (aLA) protein, ubiquitously present in milk,^5–7^ in an acidic solvent, show cyto-toxic activities against cancer cells. Such potential applications make the MG states interesting. However, the structural and functional characterization of the MG is largely lacking, since the MG state is not directly amenable to crystallization.

Intrinsic disordered proteins (IDP)^8^ also lack well-defined conformation, but can participate in functional ligand binding.^9^ The IDP inherently posses meta-stable conformations.^10^ The structures of the protein at MG state are highly dynamic and flexible.^11^ Experimental studies using the time resolved fluorescence suggest that conformation fluctuations in the molten globule state occur in nanosecond time scales.^12^ However, it is not evident that the conformation fluctuations in MG state have resemblances with those in IDP, since the MG states are induced by external conditions quite unlike the IDP.

With this backdrop, we study the microscopic nature of conformation fluctuations in MG state of aLA. Ca^2+^ ion is coordinated to this protein under physiological conditions by the carbonyl oxygen of Lysine (LYS)79 and Aspartate (ASP)84, side chain carboxylates of ASP82, ASP87, ASP88 and the crystal waters. At acidic pH, protons compete with Ca^2+^ for the carboxylate oxygen. Ca^2+^ ions can not coordinate with the protein due to uncompensated negative charge-charge interaction at the ion binding site. Fluorescence and Circular dichroism (CD) data reveals that binding of Ca^2+^ to aLA show pronounced change in structure and function of the protein.^13–15^ Calorimetric study shows that Ca^2+^ ion reduce molecular flexibility and increases its thermal stability.^16^ The removal of the ion reduces the overall stability of the protein^17^ resulting MG states of the protein. The MG state of aLA show binding with fatty acids^18–22^ like oleic acid (OA) having cyto-toxic activities. Thus, aLA in MG state acts like a carrier of cytotoxic factor. These complexes are called XAMLET, namely, aLA made lethal where X stands for the name of the mammal.^23–25^ For instance, for human milk protein, the complex is called HAMLET.

Since lowering pH is essential for the formation of the MG state of aLA, we perform biased simulations, using constant pH molecular dynamics method (CpHMD).^26^ The protonation states of the titrabale groups are updated using the Monte Carlo moves as per the generalized Born energy cost due to changes in the charged states of the residues in a continuum.^26^ Such scheme has been utilized using continuum solvent model to get MG states of aLA.^11^ MG state has been shown in canine milk lysozyme,^27^ using normal molecular dynamics simulation at different temperature and replica exchange umbrella sampling method at implicit solvent conditions. However, the conformation fluctuations in the MG state have not been addressed in these works.

We perform a hybrid scheme where explicit water molecules are taken into account in the MD simulations but the charged state updated using the implicit solvent model. The water molecules are explicitly taken to ensure coordination of the ion in neutral pH condition. We find that the structure and ion coordination from the hybrid simulation compare well to those from normal all-atom calculations at neutral pH. Next, we study aLA at low pH in an explicit solvent by using the CpHMD algorithm.^28^ We employ the dihedral angle based principal component analysis,^29^ the density based clustering and the machine learning techniques^29–35^to identify meta-stability in MG state. We also examine internal motion of the protein at neutral and pH=2.0 condition in terms of Lipari-Szabo order parameter (*S*^2^)^36^ and internal correlation time (*τ*_*e*_).^37, 38^ Our analysis reveals the presence of meta-stability via a number of helix residues in the crystal structure. These residues lack well-defined secondary structure in MG state, show smaller structural persistence (*S*_*P*_), larger dihedral auto correlation timescale and dynamic correlations with the fatty acid binding residues.

## 2 Methods

### 2.1 System preparation

The protein used in this study is Bovine *α*-lactalbumin in both holo (with Ca^2+^ ion, RCSB PDB ID: 1F6R) and apo (with Ca^2+^ ion, RCSB PDB ID: 1F6S)^39^ form. Initial structures are shows in supplementary information (SI Fig.S1). The crystal structure data of the protein tells that the protein has a *α*-helical domain and a beta sheet domain separated by a cleft. Both initial structures have six identical chains, in which we choose only the first chain for the simulation.

### 2.2 Simulation Details

We use the protocol suggested by Swails et al.^28^ to simulate CpHMD in explicit solvent. At first, MD is carried out with constant set of protonation state, assigned initially. At some point the MD is stopped, and the solvent is stripped. At this point, potential is changed to Generalized Born (GB) potential and protonation state changes for each titrable residues randomly. The electrostatic energy due to this transition and Monte Carlo decision regarding to accept or rejection of new protonation state is done the same way as in Mongan et al.^26^ If any attempt is accepted, the solute is frozen and MD is performed on the solvent to relax the solvent distribution around the new protonation states.

AMBER gives definitions for titrating side chains of aspartate(ASP), glutamate(GLU), histidine(HIS), lysine(LYS), tyrosine(TYR) and cysteine(CYS). We use ASP and GLU for titration at pH=2. Due to high acidic pH(=2), HIS is excluded. At neutral condition, pH=7, only HIS is considered for titration as *pK*_*a*_ *HIS* = 6.5. Since, CpHMD simulation in AMBER do not support C or N terminal titrable residues, they are excluded for titration during simulation.

Systems are parametrized using the AMBER FF10 force field(ff)(cite). This ff is equivalent to the AMBER FF99SB for protein. A 10Å TIP3P^40^ water is used as solvent while a truncated octahedron box has been placed around protein. The system is neutralized by placing 8 Na^+^ atoms randomly.

After making the topology, the system is minimized for 1000 steps following both steepest descent and conjugate gradient. The system is then heated at constant pressure, varying temperature linearly up to 300K. Salt concentration of the system is set at the default value of 0.15M. All bonds involving hydrogen atoms were constrained using SHAKE algorithm^41^ and temperature is maintained at 300K using Berendsen thermostat with a time constant 2ps. The solvated protein is then equilibrated for 5ns at constant temperature and pressure. Consecutive MC trials are separated by 2 femtosecond(fs) time steps. Production run is done using similar protocol for 100ns. Table.S1 shows predicted *PK*_*a*_ value of titrable residues obtained from the simulation. This prediction is considered accurate if offset value i.e. absolute value of the difference between the predicted *pK*_*a*_ and system pH is less than 2.0.^26^ Table.S1 shows that most of the titrable residues have offset value less than 2.0. All parameters are same in case of CpHMD in pH=7.

We also perform normal molecular dynamics of holo form of *α*-lactalbumin protein using AMBER FF99SB force field. All parameters are remained same as earlier CpHMD case, only difference is CpHMD is not applied in holo case.

### 2.3 Analysis

#### 2.3.1 Structural characterization

We calculate: (1) The root mean square fluctuations (RMSF) for the backbone *C*_*α*_ atoms. (2) The radius of gyration (*R*_*g*_) is calculated as average distance of *C*_*α*_ atom from their centre of mass 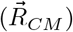. The square of *R*_*g*_ is defined as,

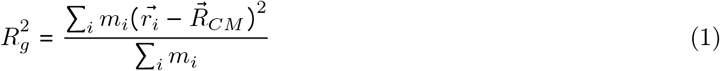

Where *m*_*i*_ and 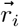 is the mass and position of ith *C*_*α*_ atom. (3) Native contacts are taken to exist between two residues if two specified atoms of the two residues are closer than a specific cutoff value. All analysis are done using CPPTRAJ tools of AMBER.^42^

#### 2.3.2 Dihedral principal component analysis

Biomolecular system like protein is very well described by backbone dihedral angle *ϕ* and *Ψ*. Hence, we use principal component analysis on dihedral angles based on newly developed dPCA+ method by Sittel et al.^29^ Traditional principal component analysis sometime produce errors in the computation of covariance matrix and projection of circular coordinates on the eigenvector. Hence, dPCA+ method minimize these errors by shifting the dihedrals periodically to set maximal sampling gap at the periodic boundary. This method mainly make use of well known fact that dihedral angles do not cover full space of dihedral space [−*π*: *π*] due to steric hindrance but are bounded in specific region. One can obtain free energy landscape by applying dPCA+ method to MD data of protein. Based on the shape of one dimensional projection of free energy landscape, further important principal components can be chosen for analysis.

#### 2.3.3 Density based clustering

We perform a robust density based geometrical cluster analysis^33^ over a landscape in the hyperspace spanned by the dihedral PCs. For every structure in the trajectory, we count the number of frames within a fixed radius R from the given frame inside the hypersphere. Normalization of the count gives density of sampling probability P. Free energy is estimated using the equation Δ*G* = −*k*_*B*_*TlnP*. At first, an energy cut off value at relatively low free energy (F < 0.1 *k*_*B*_*T*) is defined. All structure below the cut-off are considered, and all others are ignored. Selected frames that are closer than a certain lumping radius (*d*_*lump*_) will be assigned to a same cluster. The energy cut off is increased gradually at a step of 0.1 *k*_*B*_*T* until all clusters are converged at the energy barrier. In this way, all structures will get specific cluster membership. Nagel et al. shows that in most cases *d*_*lump*_ itself is a good choice for clustering radius R. Here, we consider both R and *d*_*lump*_ equal to 0.521.

#### 2.3.4 Dynamical clustering

Density based geometrical clustering is expected to give valid description of primary description of the system if conformational state of different geometrical state are separated by large barriers. But, sometime small structural change is observed in case of functional motion in large protein. Even in case of rare transition between states, geometrically different but dynamically close states are artificially separated. Meanwhile, in case of low sampling dynamically distinct states are considered as geometrically close and thus may be allotted wrongly. This error can be minimized by considering dynamic clustering method that combine MD frames which are near in time evolution instead geometrical evolution. Here, we use the most probable path (MPP) algorithm developed by Jain et al.^31, 32^for dynamical clustering. At first, given a set of microstates, the transition matrix of these states is calculated. For a given state, if self transition probability is lower than certain metastability criterion Q_*min*_ *ϵ* (0,1], then the state will be merged with the state having the highest transition probability and lower free energy. The process is reiterated until for a given Q_*min*_ no further transition happen.

#### 2.3.5 Essential internal coordinates (EC)

We have used supervised machine learning techniques to find essential coordinates for meta-stability in the MG state. We identify an essential coordinate of the system in the meta-stable state using extreme gradient boosting (XGBoost) algorithm.^43^ In this method, given a trajectory of MD coordinates in terms of dihedral angle and a meta-stable state obtained from clustering, a machine learning model is constructed by minimizing a loss function. Overall accuracy of the model can be estimated by dividing the available data into train and test set. The importance of a coordinate is given by the gain of the loss function value. Any MD coordinate which has high gain in loss function is more important for characterizing the state than others. Given a trained model, all the dihedrals are sorted by their importance. When an non-essential dihedral is discarded, the accuracy of the model does not change significantly. Thus, one can identify essential internal coordinates that are necessary to discriminate the states explicitly. Here, all XGBoost parameters are chosen as in Ref.^30^

#### 2.3.6 Structural persistence

The structural persistence (*S*_*p*_) parameter is defined as:^44^

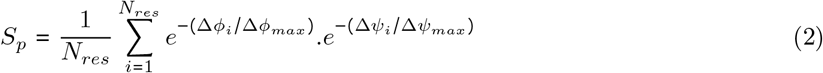

Here, *N*_*res*_ denotes total number of residues. Δ*ϕ*_*i*_ and Δ*Ψ*_*i*_ represents change in dihedral *ϕ* and *Ψ* of ith residue with respect to initial crystal structure. Δ*ϕ*_*max*_ and Δ*Ψ*_*max*_ denotes the maximum permissible change in dihedral angle in Ramachandran space without considering direction. *S*_*p*_ = 1 denotes no change in protein conformation, whereas low *S*_*p*_ represents higher deviation from the reference structure. Structural persistence for a single residue is calculated in similar manner. Although, summation and averaging over time is done for a single residue instead of the full protein.

#### Correlation functions and order parameter

In general, dipolar interaction between two nuclei can be measured using NMR. The correlation function for such cases is defined as:

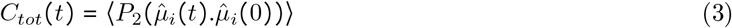

where 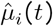is the unit dipole moment vector pointing along the given dipole, and *P*_2_ (*x*) = (1/2)(3*x*^2^ − 1) is a second order Legendre polynomial. ⟨.. ⟩ represents that averaging is performed over all given dipoles at different time origins. For macromolecules like protein, timescale associated with internal motion is faster than overall motion, henceforth these two motion could be considered independently. Hence, *C*_*tot*_ (*t*) can be factorized into correlation functions for overall motions (*C*_*O*_ (*t*)) and internal motions (*C*_*I*_ (*t*)) of the protein. *C*_*I*_ ((*t*)) can be calculated after removing center of mass of rotational and translational motion of protein with reference to initial structure.

The relaxation behavior of *C*_*I*_ (*t*) can be quantified by fitting the *C*_*I*_(*t*) decay curve with the Lipari-Szabo model free approach^36^

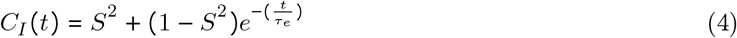

Where *S*^2^ and *τ*_e_ are defined as order parameter and effective internal correlation time of given dipole. It is important to note that *S*^2^ and ⟨*τ*_*e*_⟩ can be calculated from NMR relaxation data. Parameter *S*^2^ denotes the level of spatial rigidity of a given dipole. *S*^2^=1 denotes totally rigid state, whereas 0 value represents entirely free motion. Here we considered two dipoles of protein backbone i.e. N-H and C-N dipoles. Here, internal correlation functions (*C*_*I*_(*t*)) averaged over all C-N and N-H dipoles respectively at both neutral and pH=2. *C*_*I*_ (*t*) is calculated over the last 50ns of the production run.

#### 2.3.8 Dynamical cross correlation

Correlated motion between various protein segment can be defined in terms of simple cross correlation *C*(*i, j*)functions^45, 46^ of various residues. If Δ*r*_*i*_ and Δ*r*_*j*_ are the displacement vectors of i-th and j-th residue of *C*_*α*_ an atom of the protein, then *C*(*i, j*) is defined as

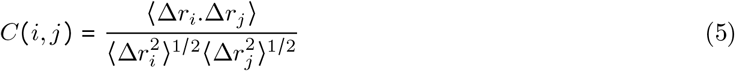

Where the angular bracket signifies ensemble average. *C*(*i, j*)varies between -1 (anti correlated motion) to +1 (correlated motion) range. Movement of correlated residues are in same direction, whereas anti correlated residues are in opposite direction. It is noted that we have used trajectory at *t* = 0 as the reference structure, and all other structures are aligned with respect to that reference structure. Calculations are done using Bio3D suite of R programming package.^47^

## 3 Results

We perform CpHMD simulation in an explicit solvent in calcium ion free (apo-) and bound (holo-) aLA at neutral medium (pH=7.0) and apo aLA at acidic medium (pH=2.0). We also carry out normal MD simulations at neutral pH on holo-aLA. At first, we validate structures obtained from CpHMD and normal MD simulation of the holo-structure in neutral pH. Supplementary information, SI Fig.S2 shows both structures overlapped, where the root-mean-square deviation between the atoms is found to be only 0.479 A^*o*^ with similar overall secondary structures. This shows the validity of the hybrid simulations. We compare constant CpHMD MG conformation of apo aLA in acidic medium to that of normal MD conformations of holo aLA in neutral solvent.

### 3.5 Conformations in the MG state

SI Fig.S3(a) shows RMSF per residue for holo aLA at neutral solvent using normal MD and apo aLA at acidic solvent using the CpHMD simulations. The data suggest that RMSF increases at acidic pH compared to the holo-alA in neutral condition. Radius of gyration (*R*_*g*_) as given in Eq.1 measures compactness of any protein. *R*_*g*_ is computed for each conformation. SI Fig.S3(b) shows histograms (*H*(*R*_*g*_)) of *R*_*g*_ for both holo-aLA in neutral and acidic pH. Average value of *R*_*g*_ obtained from simulation at MG state is 14.72 A^*o*^ is slightly larger than holo-aLA in neutral case (Average *R*_*g*_ = 14.41 A^*o*^). Qualitatively, this is in agreement to earlier experimental observations and simulations.^48–51^

The native contacts in a denatured protein decrease, as revealed from the nuclear magnetic resonance (NMR) chemical shift dispersion spectra.^11, 52, 53^ SI Fig.S3(c)-(d) shows native contact map of holo-aLA and that at pH=2 with 7 Å cut off between *C*_*α*_ atom as reference atom of the residues. The native contacts decrease at acidic pH. Non-native contact map is shown in SI Fig.S3(e) for neutral and (f) for pH=2 case. The maps show that non-native contacts, on the other hand, increase at acidic pH than in the neutral case, in agreement to earlier simulation studies.^11^

Earlier works show changes in the solvent accessible surface area (SASA) in the molten globule state.^11^ Fluorescence studies suggest that solvent accessibility of Tryptophan residue in MG state increases as compared to neutral state.^54, 55^ Earlier implicit solvent based analysis in MG state show similar result.^11^ Accordingly, we calculate the SASA. We observe increase of SASA of Tryptophan(W) at MG state. Table.1 shows the SASA values of Tryptophan(W) residue at neutral and molten state. The structural persistence (*S*_*P*_) (see Eq.2) is sensitive to changes in structural preferences. We show the joint probability distributions of SASA and *S*_*P*_ at both neutral and acidic pH in Fig.1(a) and (b) respectively. Compared to the neutral pH (Fig.1(a)) we observe that the lower structural persistence is correlated with higher SASA value at acidic pH (Fig.1(b)).

**Table 1:** SASA value of Tryptophan(W) residues at neutral and acidic pH.

| Condition | W60 | W104 | W118 |
| --- | --- | --- | --- |
| Neutral | $4\text{\AA}^2$ | $16\text{\AA}^2$ | $19\text{\AA}^2$ |
| pH2 | $19\text{\AA}^2$ | $96\text{\AA}^2$ | $34\text{\AA}^2$ |

**Figure 1:**
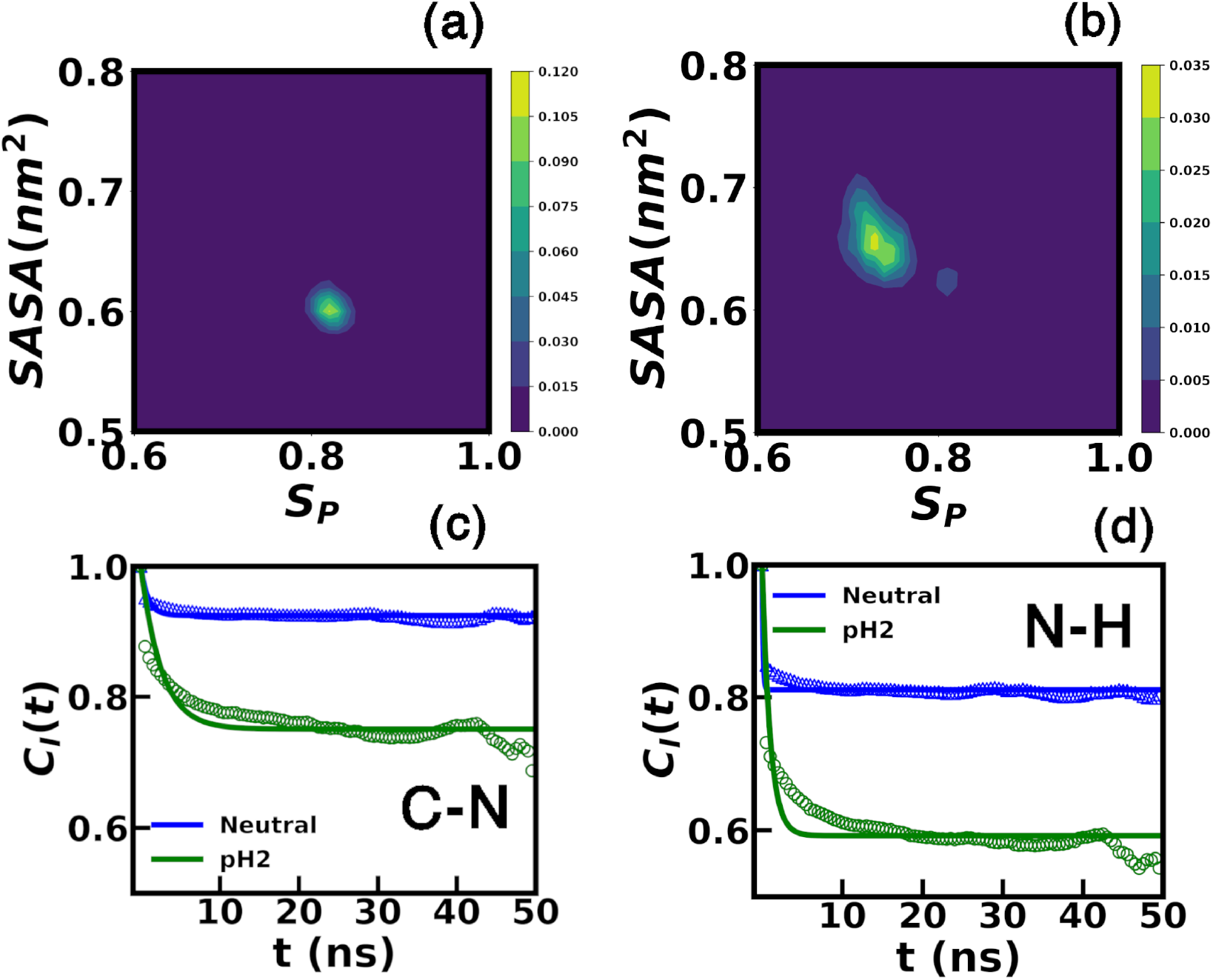
Joint probability distribution of SASA and *S*_*P*_ at (a) neutral and (b) pH2. Internal correlation functions (*C*_*I*_ *t*) for the (c) C-N bond dipole and (d) N-H bond dipole. The scatter plot shows the original curve, while the fitted line is shown in solid.

We also consider dynamics of protein backbone N-H and C-N dipoles. Fig.1(c) and (d) shows internal correlation functions (*C*_*I*_ (*t*)) (see Eq.3) averaged over all C-N and N-H dipoles respectively for alA at acidic pH and holo-alA at neutral condition. *C*_*I*_ *t* decay slowly at acidic pH as compared to neutral cases. We quantify the correlation relaxation further by fitting *C*_*I*_ (*t*) using Eq.4 and estimated the Lipari-Szabo order parameter *S*^2^. The fitted parameters of Eq.4 are in Table.2 for N-H dipoles and C-N dipoles. The data suggest that lowering the pH reduce the order parameter (*S*^2^) values for both dipole, while the correlation time (*τ*_*e*_) increases in comparison to the neutral holo-aLA.

**Table 2:** Order parameter (*S*^2^), internal correlation time (*τ*_*e*_) for the backbone N-H dipole and C-N dipole of *α*-lactalbumin protein at both neutral and pH2

| Dipole | Condition | $S^2$ | $\tau_e(\text{ns})$ |
| --- | --- | --- | --- |
| N-H | Neutral | 0.81 | 0.12 |
| N-H | pH 2 | 0.59 | 0.99 |
| C-N | Neutral | 0.92 | 0.91 |
| C-N | pH 2 | 0.75 | 2.58 |

### 3.2 Meta-stability in the MG state

Next, we consider fluctuations in the MG state in the dihedral angles using the dPCA+ method^29^ (see in the Method section). One dimensional free energy landscapes for PC1 to PC5 are shown in Fig.2(a)-(e) all of which have the presence of meta-stable states. Representative cases of other PC 6-10, where meta-stability is absent, are shown in SI Fig.S4. The density based geometrical cluster analysis^33^ over the hyperspace spanned by the dihedral PCs (see Methods for details) provide 56 micro-states. Now, we use the most probable path (MPP) algorithm (see Methods for details) to reduce micro-states into a set of meta-stable states. We obtain 27 micro-states using lag time of *τ* =40 pico second and minimum meta-stability of *Q*_*min*_=0.92, eliminating spurious transition in the vicinity of the energy barrier by coring as discussed in Nagel et al.^35^

**Figure 2:**
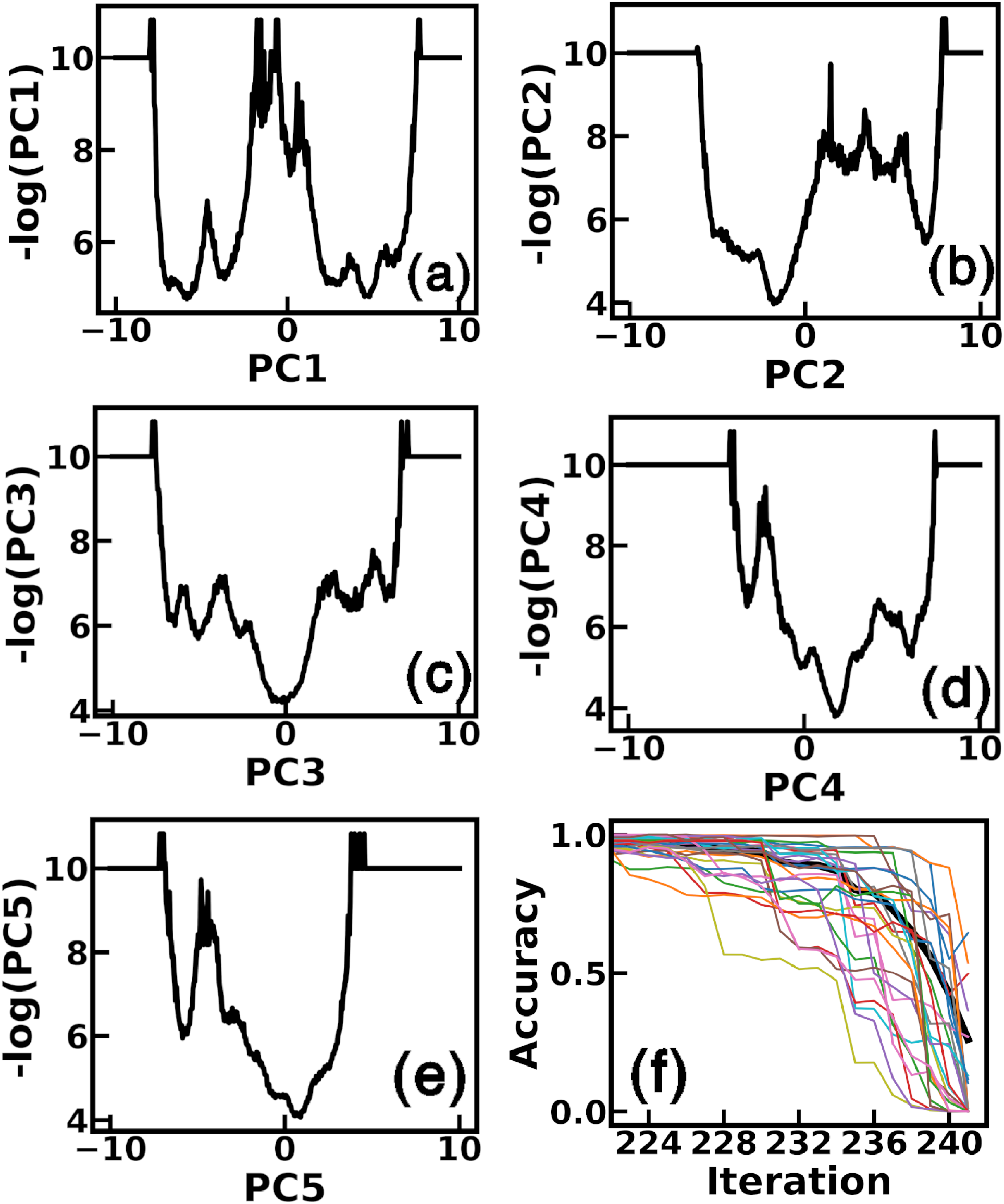
Free energy landscape obtained from dPCA+ for (a) PC1, (b)PC2, (c)PC3, (d)PC4, and (e)PC5. (f) Accuracy loss plot of XGBoost classifier. Figure is shown as a function of number of discarded coordinate. Accuracy of all metastable states drops drastically upon removing of last 10 coordinate.

Given the meta-stable states, we use the XGBoost model (see in the Method section) on full MD data to identify the relative importance of the dihedral angles in the MG state. Fig.2(f) shows an accuracy plot of XGBoost algorithm. The thick black line represents state averaged accuracy. The accuracy remains constant, discarding up to 232 coordinates. Accuracy decreases sharply for most of the states by discarding the last 10 coordinates. These coordinates, shown in Table.3 as per decreasing importance with the secondary structure element in the crystal structure, are the essential coordinates (EC). We highlight in red these residues over the crystal structure (Fig.3(a)). The most essential coordinate is *Ψ*80 as it is obtained at the final iteration of the XGBoost, removing all other coordinates. We find that only *ϕ*122 and *ϕ*114 belongs to loop residues, while all other belongs to helix residues. The essential coordinate *Ψ*80 belongs to amino acid residue PHE80, which belongs to the helix near the calcium binding loop. Similarly, the residue LYS79 having the essential coordinate *ϕ*79 directly coordinates to calciumion.

**Table 3:** Residues list which posses EC obtained using XGBoost algorithm. Secondary structure for each residue in initial crystal structure is mentioned.

| EC | Residue Name | Crystal structure |
| --- | --- | --- |
| $\psi 80$ | Phenylalanine(PHE) | Helix |
| $\psi 18$ | Tyrosine(TYR) | Helix |
| $\phi 79$ | Lysine(ASP) | Helix |
| $\phi 17$ | Glycine(GLY) | Helix |
| $\psi 109$ | Alanine(ALA) | Helix |
| $\phi 122$ | Lysine(LYS) | Loop |
| $\phi 114$ | Lysine(LYS) | Loop |
| $\psi 99$ | Valine(VAL) | Helix |
| $\psi 105$ | Leucine(LEU) | Helix |
| $\phi 77$ | Cysteine(CYS) | Helix |

**Figure 3:**
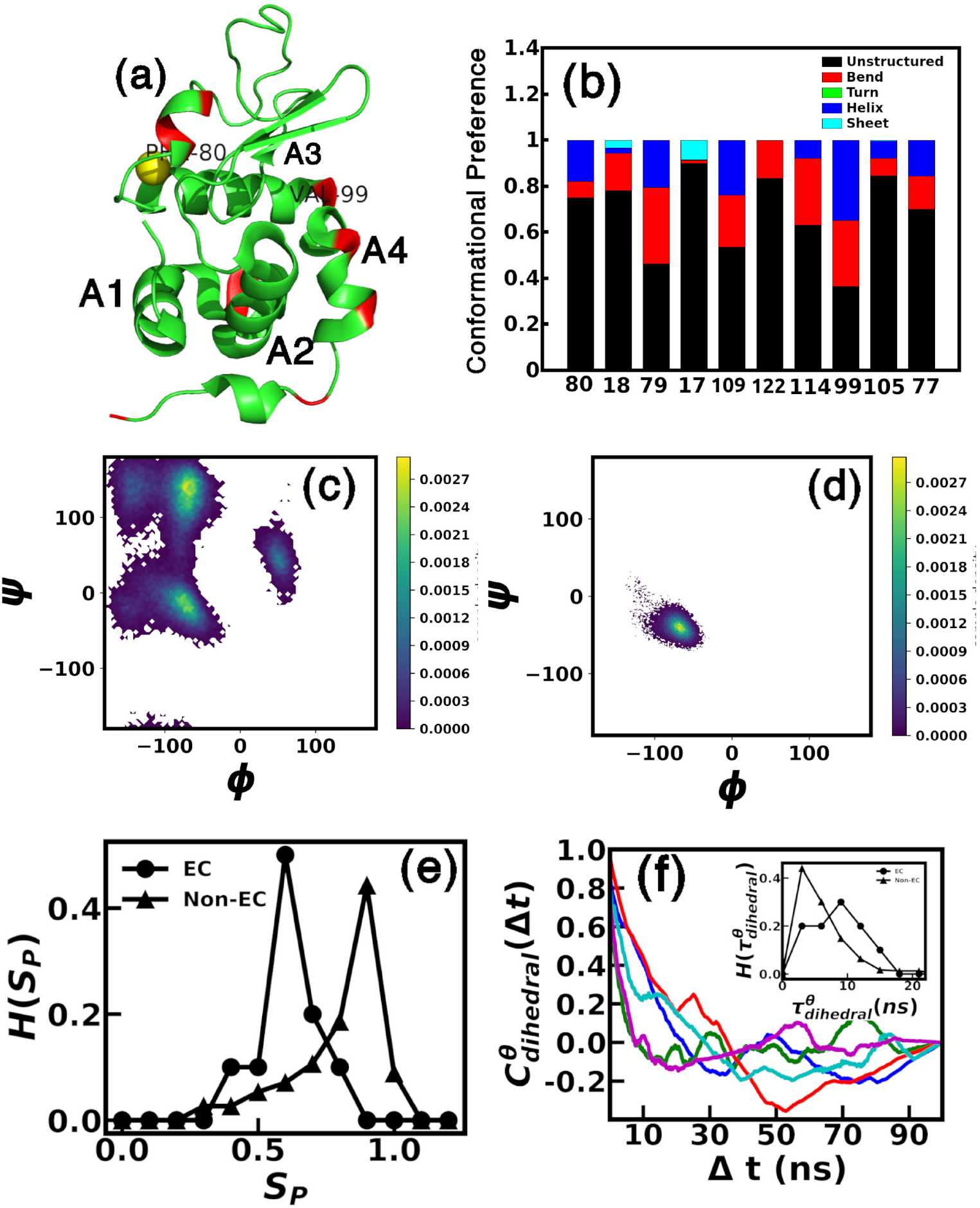
(a) Initial structure of *α*-Lactalbumin. Essential coordinates are marked in red color. (b) Conformational preference of residues having essential coordinates. *ϕ*-*Ψ* correlation plot for residue (c) Phenylalanine80 and (d)Valine8. (e) Histogram of structural persistence (*S*_*P*_) for residues containing essential coordinates and non-essential coordinates. Residues having essential coordinates have less structural persistence. (f) Dihedral auto correlation function of first 5 essential coordinates (Blue:*Ψ*80 of Phenylalanine, Green:*Ψ*18 of Tyrosine, Red:*ϕ*79 of Lysine, Cyan:*ϕ*17 of Glycine, Magenta:*Ψ*109 of Alanine). Inset shows histogram of correlation timescales of essential and non-essential coordinates.

We find in Fig.3(b) that most residues having the essential coordinates prefer unstructured or bend conformation, suggesting that these residues lack well-defined secondary structural element in the MG state. Next, we show the *ϕ* − *Ψ* correlation plot for residue PHE80 in Fig.3(c). Comparing to the Ramachandran plot, we find that the conformations in PHE80 mostly lie within helix, sheet and unstructured region. This is consistent to the data in Fig.3(b). On the other hand, the *ϕ* − *Ψ* correlation plot for VAL8 which is assigned as non-essential coordinate, suggests conformations only confined within the helix region (Fig.3(d)). Next, we characterize essential and non-essential coordinates in terms of *S*_*P*_. We consider two data sets, one of the residues having the ECs and the other consisting of the remaining residues. The histograms of the structural persistence (*H(S*_*P*_)) for these sets are shown in Fig.3(e). We find that residues containing EC coordinate posses low *S*_*P*_, whereas non-EC coordinates posses higher value of

Next, we compute the time dependent correlation functions 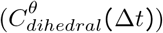 of dihedral fluctu-ations, using the method discussed in Ref.^56^ Fig.3(f) shows dihedral auto-correlation functions of first 5 essential coordinates. The TDCF is normalized by Δ*t* = 0 value. We fit the initial decay data with an exponential form 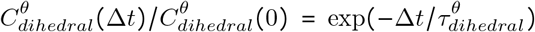. 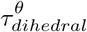 is the decay timescale. The histogram of the correlation decay timescales 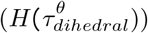 for the EC and non-EC dihedrals are shown in the inset of Fig.3(f). We observe that ECs have higher value of correlation timescales compared to non-EC. Average correlation timescales, 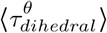 for ECs is 6.65 ns and for non-ECs 4.42 ns, suggesting that the fluctuations in the EC are longer lived than those in non-EC degrees of freedom.

Experimental data suggest that the MG in aLA is realized in absence of the Ca^2+^ ion.^17^ This is supported by our data as well. We perform CpHMD simulation with apo-aLA at neutral conditions. SI Fig.S5(a) shows dihedral PCs obtained in this case. High meta-stability is present where many of the ECs belong to stable structure in initial crystal structure (Table S2) as in the MG state. We contrast the conformation fluctuations in holo-aLA in the neutral solvent conditions. The free energy profiles obtained from the dPCA+ analysis (SI Fig.S6 (a)) show the presence of meta-stable states. However, in contrast to apo-aLA, the essential dihedral angles in the neutral case for the holo-protein mostly lie on the loop region in crystal structure (Table S3) and prefers unstructured conformation over the trajectory (SI Fig.S6). Thus, the loss of Ca^2+^ ion is the primary factor to realize the MG state of aLA.

### 3.3 Implication for functionality

It is known that in the MG state, alA bind to fatty acids, like oleic acid (OA) with negatively charged carboxylate (COO^−^) head groups and long hydrophobic tail to form cytotoxic complex.^57–59^ Experiments^59^ suggest that recognized binding sites for OA binding lies between A1 and A2 helices (Fig.3(a)) and cleft region. Table S4 shows putative binding residues of OA with initial structural element. GLY17, VAL99 and other neighboring hydrophobic residues are the probable residues participating in OA binding. Among the ECs, VAL99 of the cleft region and GLY17 of the A1 helix are putative binding residues. Thus, some residues directly participating in the OA binding bear the essential coordinates.

We relate the ECs to the SASA of the binding residues. Accordingly, we draw correlation plot between EC fluctuations with SASA values of the putative binding residues. As ligand binding is mainly due to the hydrophobic interaction, we consider hydrophobic residues of cleft region. Fig.4(a) shows a correlation plot between SASA of ILE89 and dihedral *Ψ* fluctuations of PHE80 (Fig.4(a)). Correlation plot shows different clusters corresponding to different structural element of PHE80. We also check correlation plot between SASA of ILE89 and dihedral *ϕ* fluctuations of LYS79 which play essential role in Ca^2+^ coordination (SI Fig.S7). Correlation plot shows different cluster at two different regions of conformational space which is similar as PHE80. It may be noted that the PHE80 and LYS79 are distant from ILE89, suggesting allostery^60–62^ between these residues. The correlation plots for other ECs and putative binding residue shows different clusters as the conformations of EC changes. We contrast the SASA of ILE89 with dihedral *ϕ* (Fig.4(b)) and *Ψ* (Fig.4(c)) fluctuations of VAL8 which has only non-essential coordinates. The correlation plot shows that the SASA fluctuations are not correlated to those of the dihedrals.

**Figure 4:**
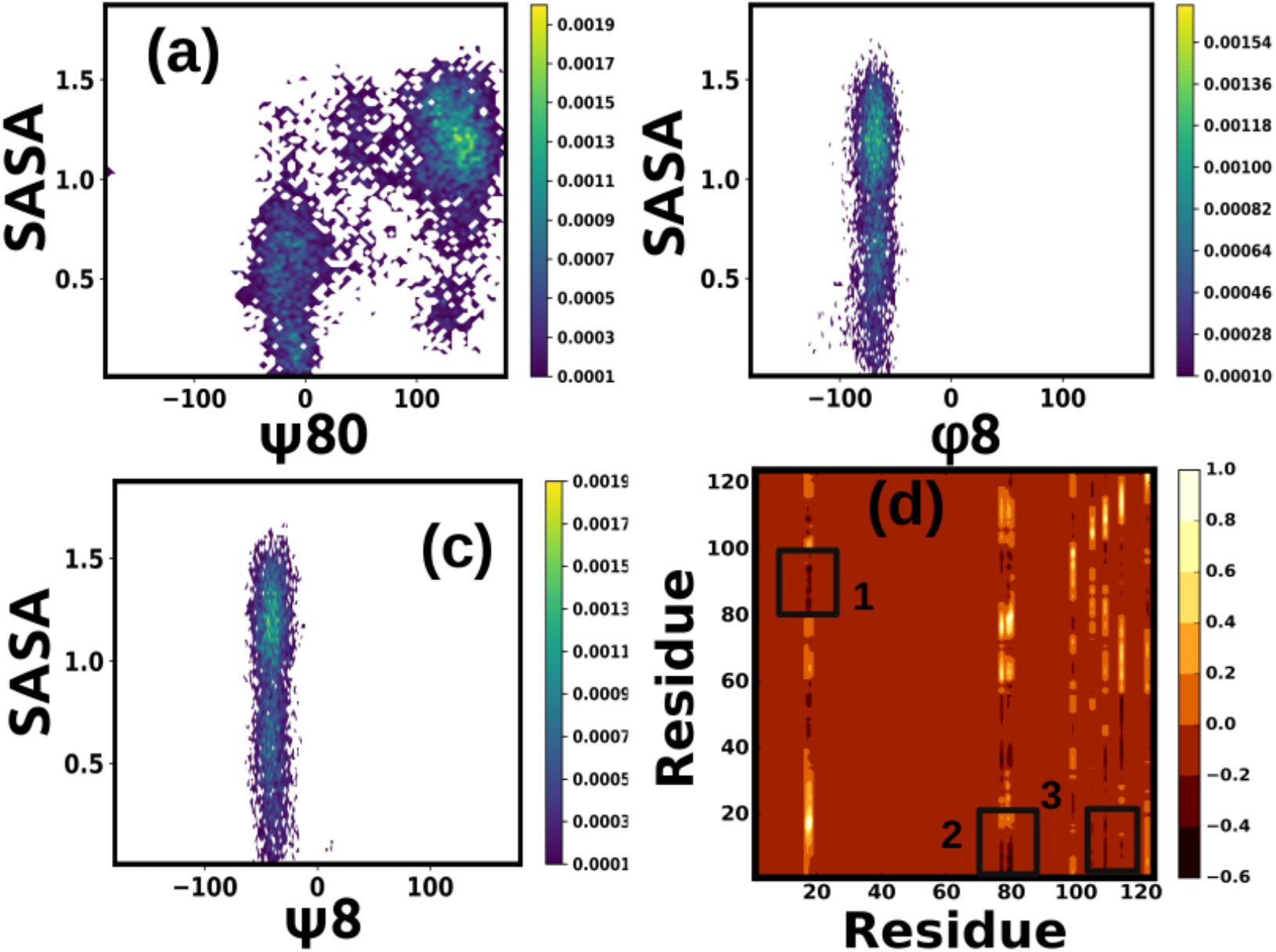
Correlation plot between SASA value of Isoleucine89 with (a) dihedral *Ψ* fluctuations of Phenylalanine80, (b) *ϕ* of Valine8 and (c) *Ψ* of Valine8. (d)DCCM map between residues having essential coordinates with all other residues. Box 1 represents DCCM map between GLY17 and TYR18 with putative binding residues of interfacial cleft, box 2 represents DCCM map between LYS79 and PHE80 with putative binding residues of A1 helix, box 3 represents DCCM map between LEU105, ALA109, LYS114 with residues of A1 and A2 residues.

We further elucidate the connection between putative binding residues and the EC via the dynamical cross correlation (Eq.5) map of *C*_*α*_ atom fluctuations belonging to these residues (Fig.4(d)). DCCM plot shows that GLY17 and TYR18 are anti-correlated with putative binding residues of inter-facial cleft (marked as 1). Similarly, LYS79 and PHE80 are dynamically anti-correlated with residues of A1 helix (marked as 2). Residues LEU105, ALA109 and LYS114 show dynamic anti-correlation with residues of A1 and A2 helices (marked as 3). Such anti-correlated motion between residues result in opening of the inter-facial cleft required for OA binding to aLA in acidic medium.

## 4 Conclusions

In conclusion, we explore the conformation fluctuations and meta-stability in MG state of aLA in terms of essential coordinate using density based clustering and machine learning approach. We find high metastability in free energy landscape of apo aLA at MG state. ECs responsible for metastability in the MG state prefer unstructured or bend conformations, although in crystal structure they possess stable secondary structure. Residues participating in coordination of Ca^2+^ ion (PHE80 and LYS79) acts as essential coordinates of the system. Thus, the removal of Ca^2+^ initiates metastability in the protein upon lowering of pH. The ECs play a major role in opening up the putative fatty acid binding sites. These features in the MG state are similar to those of IDP. Thus, the MG state can be viewed as induced disordered protein. It will be worthwhile to understand the binding of fatty acid in such disordered state of protein. Our study will be helpful to understand functionality of a protein in partly denatured conditions, as in the MG state.

## Supporting information

Suppplementary File

## 6 Acknowledgements

AGM thanks DST India, for an INSPIRE fellowship (IF170961) to conduct the research. The authors thank the Thematic Unit of Excellence (TUE) and the Technical Research Centre (TRC) at S. N. Bose National Centre For Basic Sciences for computational facilities.

