## Supplementary material for "Conformational fluctuations in molten globule state of *α*-lactalbumin": Suppplementary File

### Electronic Supplementary Information article "Conformational fluctuations in molten globule state of $\alpha$-lactalbumin"

Abhik Ghosh Moulick[a] and J. Chakrabarti[a]

[a]Department of Chemical, Biological and Macro Molecular Sciences, S. N. Bose National Centre For Basic Sciecnes, Block JD, Sector III, Salt Lake, Kolkata-700098, India

Table S1: Predicted pKa values of titrable residues during simulations. The offset value is defined as the difference between predicted pKa and system pH. Fraction of time titrable residues remain protonated during simulations.

| Residue Name | Residue | Predicted pKa Value | Offset Value | Fraction of protonation |
|---|---|---|---|---|
| 7 | GLU | 3.675 | 1.675 | 0.956 |
| 11 | GLU | 3.334 | 1.334 | 0.987 |
| 14 | GLU | 3.869 | 1.869 | 0.794 |
| 25 | GLU | 2.586 | 0.586 | 0.998 |
| 37 | ASP | 3.239 | 1.239 | 0.945 |
| 46 | ASP | 3.652 | 1.652 | 0.978 |
| 49 | GLU | 3.933 | 1.933 | 0.988 |
| 63 | ASP | 2.472 | 0.472 | 0.748 |
| 64 | ASP | 3.270 | 1.270 | 0.949 |
| 78 | ASP | 2.714 | 0.714 | 0.838 |
| 82 | ASP | 2.594 | 0.594 | 0.797 |
| 83 | ASP | 3.077 | 1.077 | 0.923 |
| 84 | ASP | 2.784 | 0.784 | 0.859 |
| 87 | ASP | 2.052 | 0.052 | 0.530 |
| 88 | ASP | 3.499 | 1.499 | 0.969 |
| 97 | ASP | 3.392 | 1.392 | 0.961 |
| 113 | GLU | 4.661 | 2.661 | 0.998 |
| 116 | ASP | 3.282 | 1.282 | 0.950 |
| 121 | GLU | 4.091 | 2.091 | 0.992 |

Table S2: Obtained EC at neutral condition, using CpHMD with apo structure in explicit solvent

| EC | Initial structure |
|---|---|
| $\phi$18 | Helix |
| $\psi$68 | Loop |
| $\psi$122 | Loop |
| $\phi$19 | Helix |
| $\phi$45 | Loop |
| $\phi$121 | Loop |
| $\phi$17 | Helix |
| $\phi$20 | Loop |
| $\psi$18 | Helix |
| $\psi$16 | Helix |

Table S3: Obtained EC at neutral condition using normal molecular dynamics simulations, with holo structure in explicit solvent

| EC | Initial structure |
|---|---|
| $\phi$68 | Loop |
| $\psi$121 | Loop |
| $\psi$120 | Loop |
| $\psi$16 | Helix |
| $\phi$63 | Loop |
| $\psi$67 | Loop |
| $\psi$66 | Loop |
| $\phi$121 | Loop |
| $\psi$68 | Loop |
| $\psi$104 | Helix |

Table S4: Putative binding sites of Oleic acid (OA) with nature and secondary element in crystal structure.

| Residue Number | Residue | Initial Structure | Nature |
|---|---|---|---|
| 8 | VAL | Helix | Hydrophobic |
| 9 | PHE | Helix | Hydrophobic |
| 10 | ARG | Helix | Basic |
| 12 | LEU | Helix | Hydrophobic |
| 13 | LYS | Helix | Basic |
| 16 | LYS | Helix | Basic |
| 17 | GLY | Helix | Hydrophobic |
| 19 | GLY | Helix | Hydrophobic |
| 20 | GLY | Helix | Hydrophobic |
| 21 | VAL | Helix | Hydrophobic |
| 23 | LEU | Helix | Hydrophobic |
| 26 | TRP | Helix | Hydrophobic |
| 27 | VAL | Helix | Hydrophobic |
| 31 | PHE | Helix | Hydrophobic |
| 32 | HIS | Helix | Basic |
| 51 | GLY | Loop | Hydrophobic |
| 52 | LEU | Loop | Hydrophobic |
| 53 | PHE | Loop | Hydrophobic |
| 55 | ILE | Sheet | Hydrophobic |
| 58 | LYS | Loop | Basic |
| 59 | ILE | Loop | Hydrophobic |
| 60 | TRP | Loop | Hydrophobic |
| 89 | ILE | Helix | Hydrophobic |
| 90 | MET | Helix | Hydrophobic |
| 92 | VAL | Helix | Hydrophobic |
| 93 | LYS | Helix | Basic |
| 94 | LYS | Helix | Basic |
| 95 | ILE | Helix | Hydrophobic |
| 96 | LEU | Helix | Hydrophobic |
| 98 | LYS | Helix | Basic |
| 99 | VAL | Helix | Hydrophobic |
| 100 | GLY | Helix | Hydrophobic |
| 101 | ILE | Helix | Hydrophobic |
| 104 | TRP | Helix | Hydrophobic |

(a)

(b)

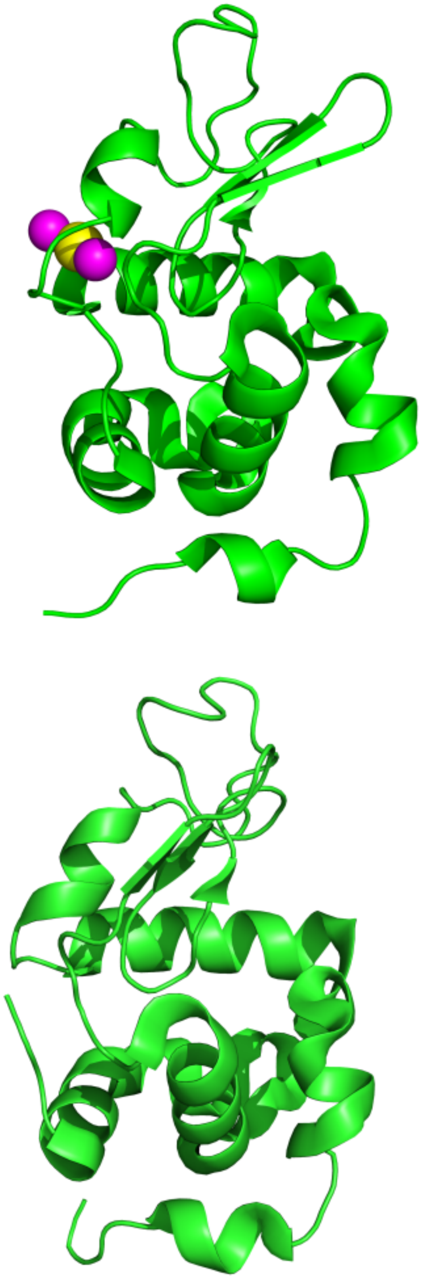

Figure S1: Initial crystal structure of $\alpha$-lactalbumin protein. (a) Holo (with $Ca^{2+}$ ion). $Ca^{2+}$ ion is shown in yellow color. Crystal waters which help in $Ca^{2+}$ coordination are in magenta color. (b) Apo structure of protein (without $Ca^{2+}$ ion).

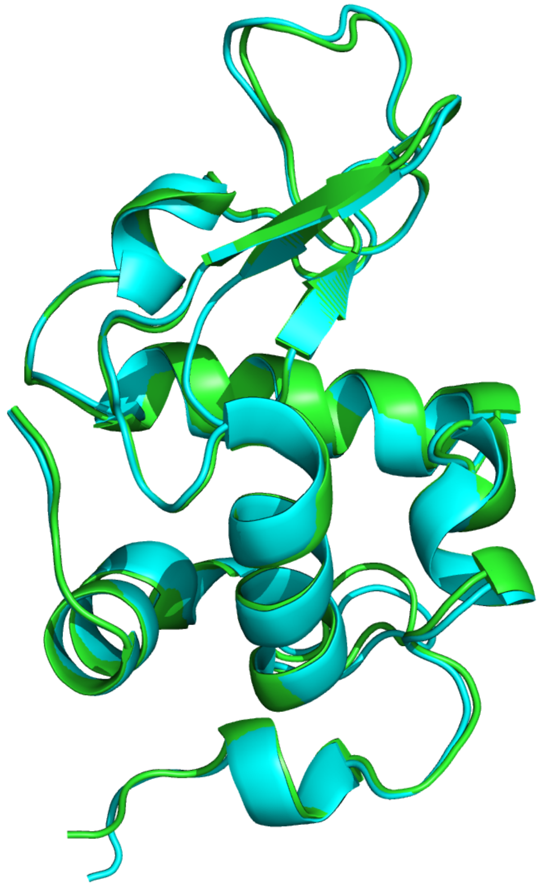

Figure S2: Overlapped average structure obtained from normal MD simulation (green) and constant pH MD simulations (cyan) at neutral pH. Root mean square distance between two structure is 0.479 A$^o$.

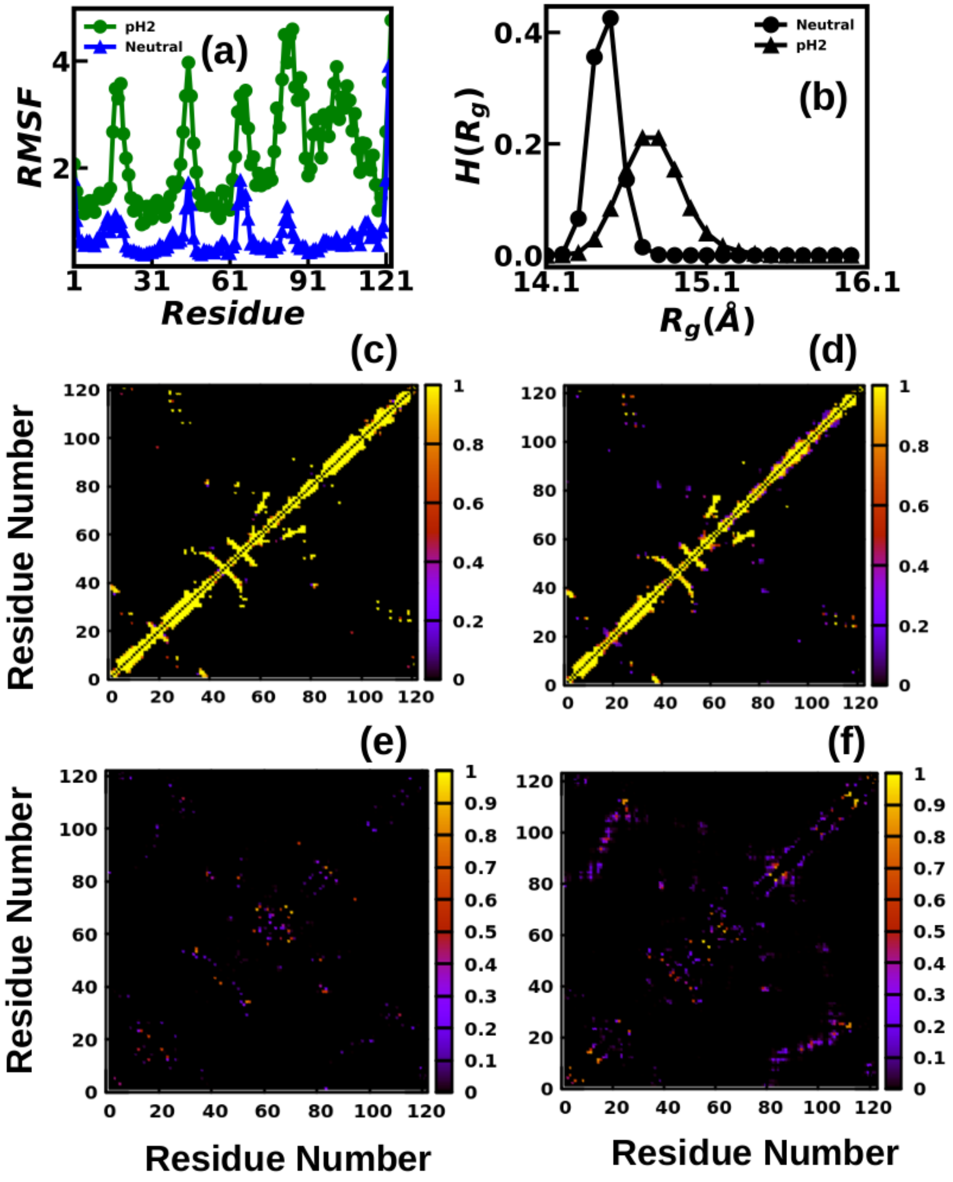


Figure S3: (a) Per residue RMSF fluctuations at both neutral and acidic pH. Histogram of, (b) radius of gyration ($H(R_g)$), Contact map of protein-native contact at (c) neutral, (d) pH2 and non-native contact at (e) neutral and (f) pH2. Native contact decrease and nonnative contact increase at pH2 as compared to neutral.

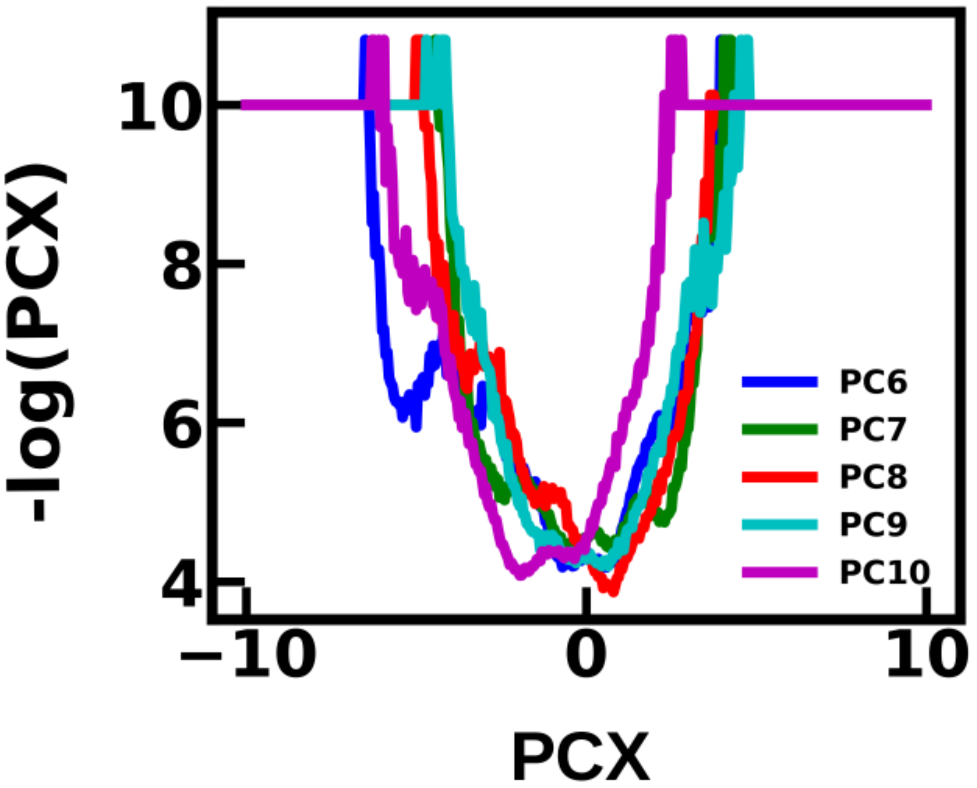


Figure S4: Free energy landscape obtained from dPCA+ for PC6-PC10. Metastability decrease gradually for higher PCs.

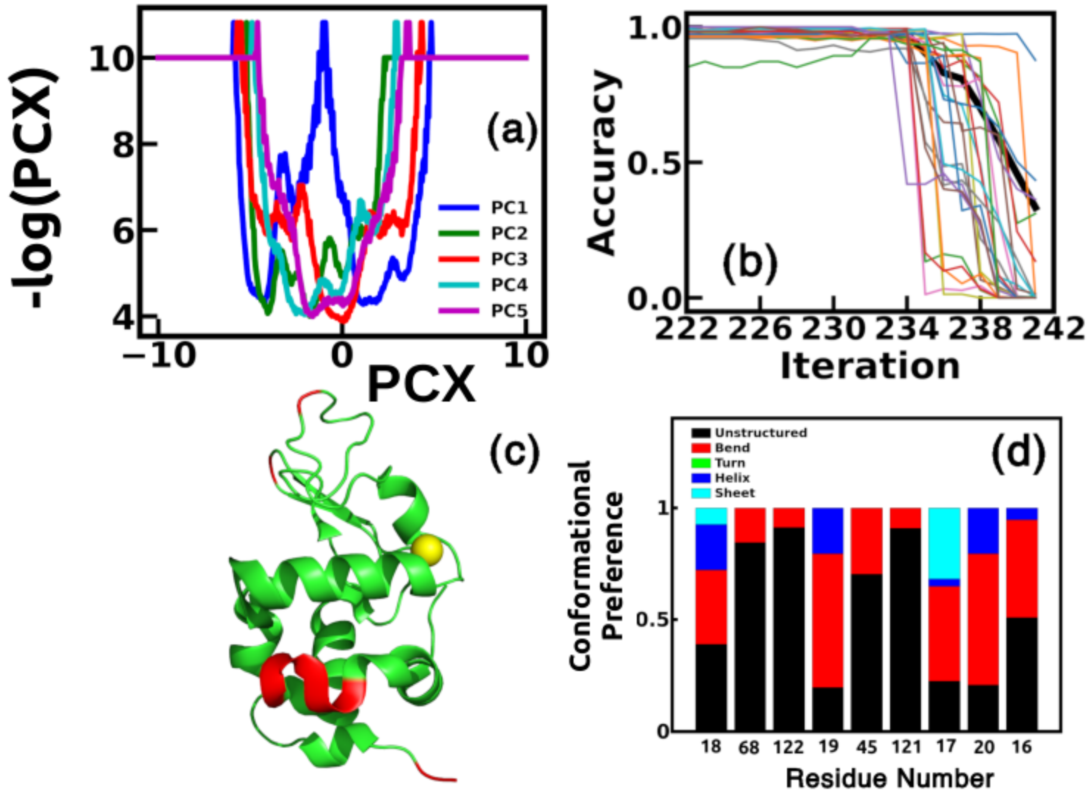


Figure S5: Identification of essential coordinate for apo ($Ca^{2+}$ ion is shown for better understanding) protein applying constant pH simulation at pH7.(a) Principal component obtained from dPCA+ analysis. PCs 1-5 are shown in figure. (b) Accuracy loss plot of XGBoost classifier. The figure is shown as a function of number of discarded coordinate. Accuracy of all metastable states drops drastically upon removing of mostly last 10 coordinates., (c) Residues having essential coordinates are marked in initial crystal structure. They are colored in red. For better understanding,k we keep the calcium ion. (d) Conformational preference of those residues having essential coordinates.

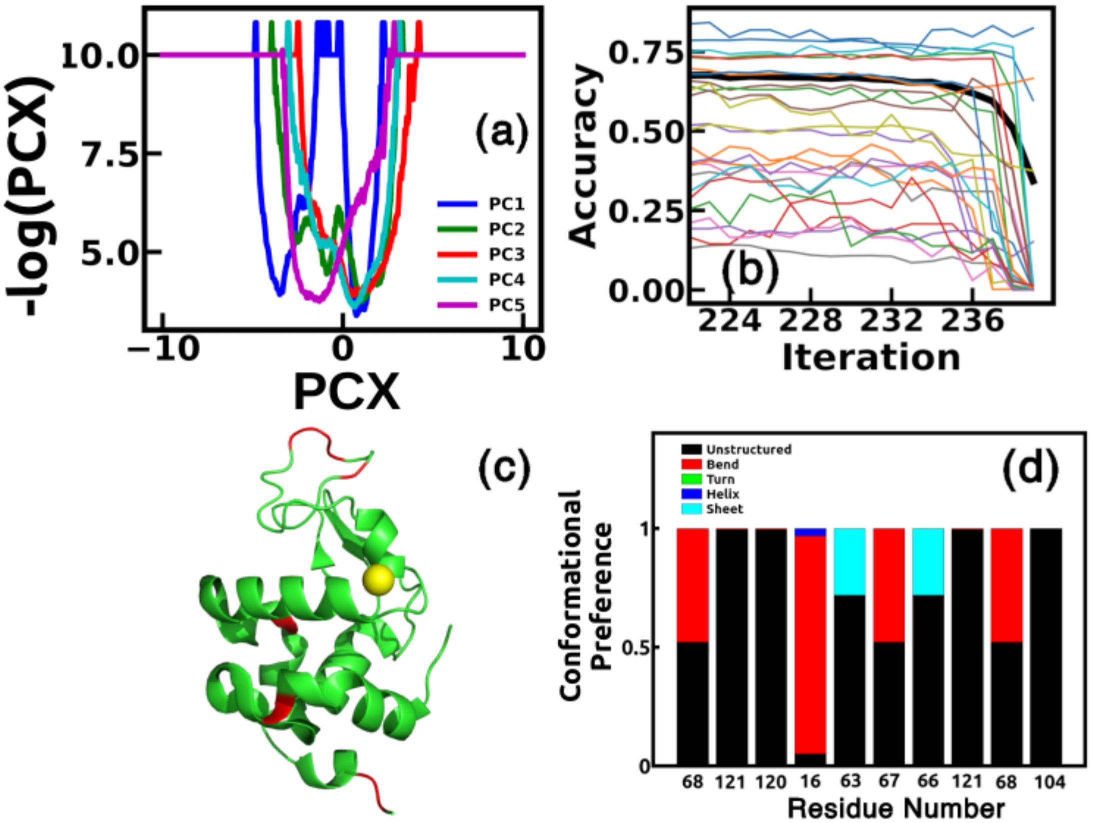


Figure S6: Identification of essential coordinate for holo protein using unbiased molecular dynamics simulation at neutral pH. (a) Principal component obtained from dPCA+ analysis. PCs 1-5 are shown in figure. (b) Accuracy loss plot of XGBoost classifier. The figure is shown as a function of number of discarded coordinate. Accuracy of all metastable states drops drastically upon removing of mostly last 10 coordinates., (c) Residues having essential coordinates are marked in initial crystal structure. They are colored in red. All non-essential residues belong to loop region. For better understanding, we keep the calcium ion. (d) Conformational preference of those residues having essential coordinates.

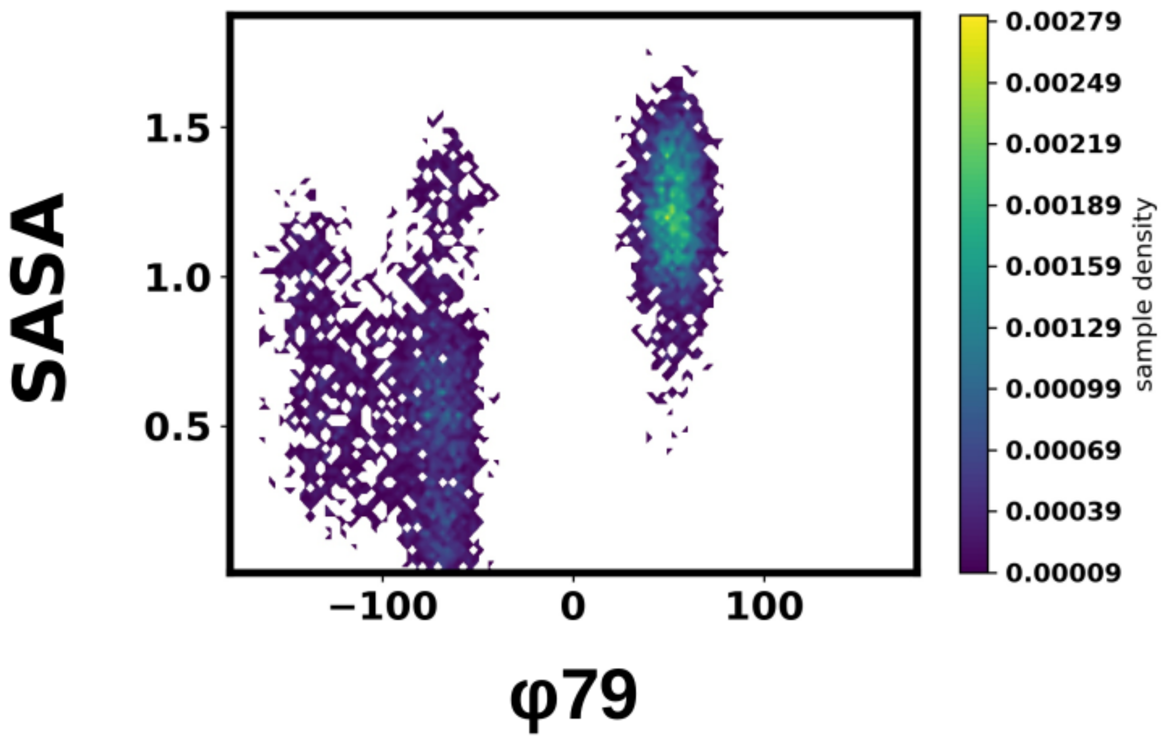


Figure S7: Correlation plot between SASA value of ILE89 with dihedral $\phi$ fluctuations of res LYS79.
